# Overcoming segmentation bottleneck in brain tissue for spatial proteomic inference at population scale

**DOI:** 10.64898/2026.10.22.761199

**Authors:** Liang You, Jamie L. Durbin, Guanqun Meng, Wayne Stallaert, Guang Li, Shikhar Uttam, Hao Feng, Jill R. Glausier, Jiebiao Wang

**Author notes:** Corresponding authors. (J.R.G.); (J.W.).

## Abstract

Tissue imaging depends on cell segmentation, whose errors carry into downstream analyses. In human brain, pretrained models struggle with complex cell bodies, and the segmentation accuracy has rarely been evaluated. Here, we introduce SPInfer, a workflow developed using thousands of expert-annotated cell bodies from 120 immunofluorescence images of human prefrontal cortex. After benchmarking eight methods, we fine-tuned the best-performing foundation model, thereby more than doubling cell segmentation accuracy. The resulting cell compositions agreed with single-nucleus RNA-seq and spatial transcriptomics. Across 14,765 images from 69 donors, zero-shot masks showed no significant association with Alzheimer’s disease pathology, whereas fine-tuned masks identified four; three of these linked OLIG2-positive cell fractions to tangle and plaque burden. Fine-tuned masks also enabled cell typing from the nuclear stain alone and detected genes associated with cell morphology. This work provides a general pipeline to segment complex tissues, enabling biological discovery in spatial proteomics at the population scale.

---

Multiplexed immunofluorescence (IF) and related spatial-omics platforms can preserve tissue architecture while measuring molecular signals in situ. This shifts spatial proteomics toward cohort-scale studies spanning dozens of donors and tens of thousands of images (*1–3*). At this scale, cell segmentation is a prerequisite for quantitative analysis. It assigns protein signals to individual cells, a step that must occur before cell type composition, morphology, or spatial organization can be measured (*4*). Downstream biological conclusions therefore inherit the quality of the segmentation masks. Errors in cell-boundary delineation could distort estimated cell fractions and neighborhood features and propagate directly into disease-association analyses (*5, 6*).

This dependency poses a particular challenge in human brain cohorts (*7–9*). At the cellular level, DAPI-stained nuclei and immunolabeled cell bodies differ within and across neuronal and glial cell populations by morphology, abundance and marker labeling. Radiating processes can make cell body boundaries ambiguous, and marker-positive processes commonly overlie the nuclei or cell bodies of other cell types, resulting in erroneous apparent co-localization. At the tissue level, cortical gray matter is organized into layers of varying cell size and packing density, and laminar cytoarchitecture differs across cortical regions (*10, 11*). White matter, by contrast, consists largely of myelinated axon bundles lined by rows of oligodendrocytes and contains few neurons (*12*). AD pathology adds further variation, including reactive gliosis, neuronal loss, and myelin and oligodendrocyte disruption (*13–16*). Annotation amid this complexity requires neuroanatomical expertise and substantial time, leaving ground truth for benchmarking and fine-tuning segmentation methods scarce.

To handle complex tissue images, cell segmentation methods have advanced from shape-prior networks such as StarDist (*17*) and SplineDist (*18*) through generalist and tissue-focused models, including Cellpose (*19*), Mesmer (*20*), InstanSeg (*21*), LACSS (*22*), and Omnipose (*23*), to foundation models such as the Segment Anything Model (SAM) (*24*) and Cellpose-SAM (*25*). Benchmarking studies show that model performance depends on tissue type and imaging modality, annotation protocol, and evaluation metric (*26–28*). However, existing evaluations have focused largely on nuclear segmentation or non-neural tissues, leaving marker-positive cell-body segmentation in postmortem human brain underexplored. Moreover, prior benchmarks treat segmentation as an isolated preprocessing step that ends at mask prediction accuracy, rarely testing whether segmentation errors alter downstream biological inference.

Here we introduce SPInfer (Spatial-Proteomics Inference) to address these gaps (Fig. 1). We generated expert annotations and benchmarked eight segmentation methods spanning classical image processing, task-specific convolutional networks, and foundation models. We then fine-tuned the most competitive candidate and compared the resulting cell compositions with single-nucleus RNA-seq (snRNA-seq) data from the same donors. We further validated SPInfer with Visium spatial transcriptomics from an independent cohort in which IF and transcriptomic data were acquired on the same sections. Applied to 14,765 prefrontal cortex images from 69 donors, SPInfer segmented 8.4 million nuclei and 557,301 marker-positive cells. In downstream analyses using identical statistical models, fine-tuned masks detected four significant associations between AD pathology and IF-derived cell fractions that the zero-shot pipeline missed. We further show that nuclear neighborhoods and graph attention networks improve spatial cell type classification beyond nuclear features alone, and that cell-body morphology covaries with gene expression in snRNA-seq data from the same donors. Together, SPInfer links fine-tuned cell segmentation to reproducible inference from spatial proteomic data.

**Figure 1:**
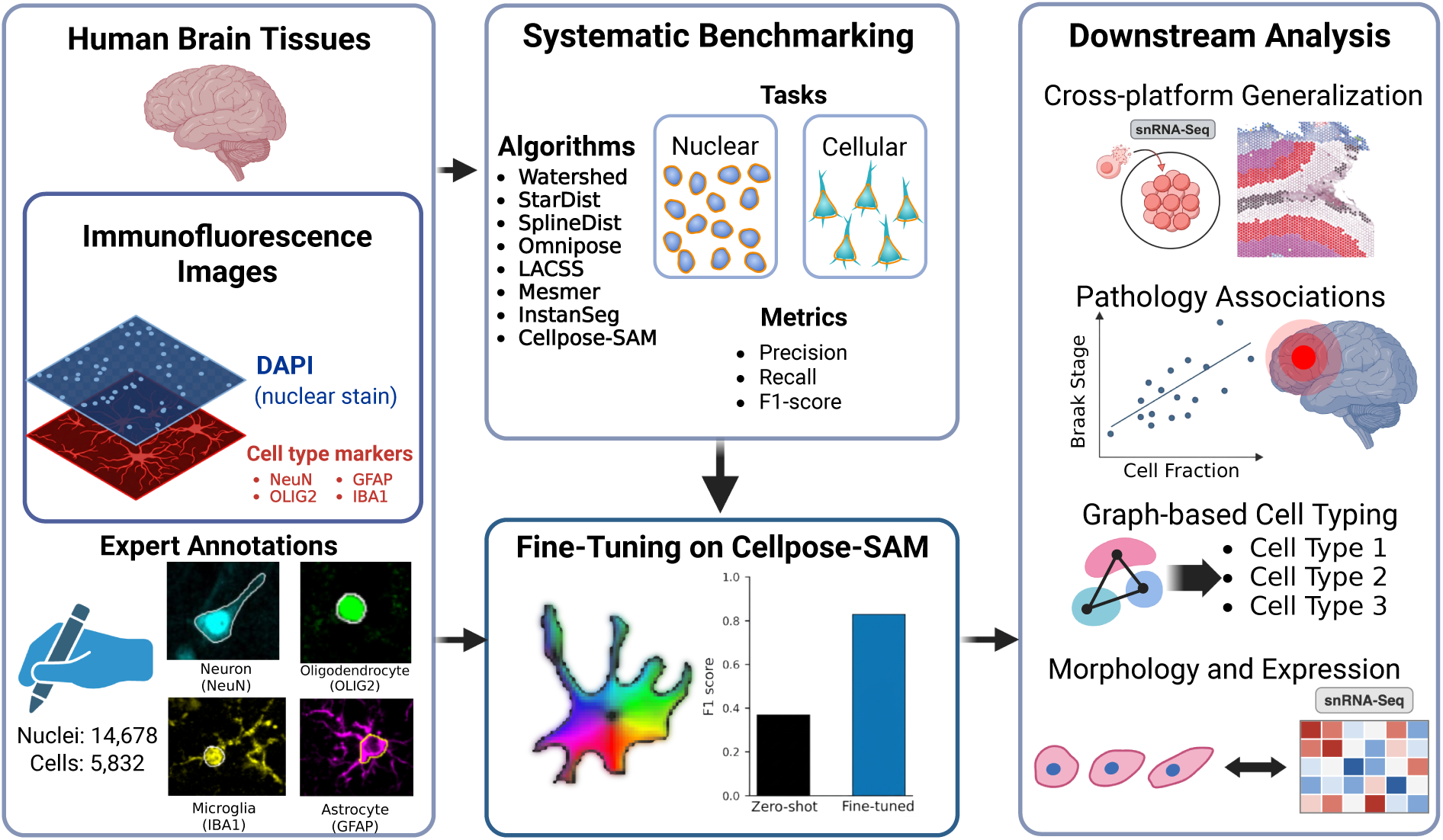
Overview of SPInfer. Left, immunofluorescence (IF) images of human prefrontal cortex pair a DAPI nuclear channel with one cell type marker (NeuN, OLIG2, GFAP, or IBA1), and experts annotated 14,678 nuclei and 5,832 marker-positive cell bodies. Middle, eight segmentation methods were benchmarked on nuclear and cell segmentation, and Cellpose-SAM was fine-tuned with the expert annotations. Right, downstream analyses. IF-derived cell fractions were compared with snRNA-seq data from the same donors and with Visium spatial transcriptomics from an independent cohort. The fine-tuned masks were then used to test associations with AD pathology, to classify cell types with a graph attention network on nuclear neighborhood graphs, and to relate cell-body morphology to gene expression.

## Benchmarking reveals the difficulty of cell-body segmentation

We first generated an expert-annotated resource of brain IF images. Each annotated image contained one DAPI nuclear channel and one cell type-specific marker channel: GFAP (astrocytes), IBA1 (microglia), NeuN (neurons), or OLIG2 (oligodendrocytes) (*8*). In total, we annotated 120 marker-labeled digital micrographs (30 per marker) containing 5,832 marker-positive cell bodies, and 30 nuclear images containing 14,678 nuclei. Annotations were generated by researchers with experience in histopathology and IF imaging and were blind to all subject characteristics.

We then benchmarked eight segmentation methods spanning classical image processing, task-specific convolutional networks, and foundation models on two tasks: nuclear segmentation and cell-body segmentation (see Materials and Methods). Performance was summarized as the F1 score, the harmonic mean of precision and recall for predicted objects matched to annotations at an intersection-over-union (IoU) threshold of 0.5. For nuclear segmentation, each method was given only the DAPI channel as input. Several methods performed well: Cellpose-SAM achieved the highest median F1 (0.91), followed by StarDist (0.88), InstanSeg (0.86), and SplineDist (0.86) (Fig. 2A).

**Figure 2:**
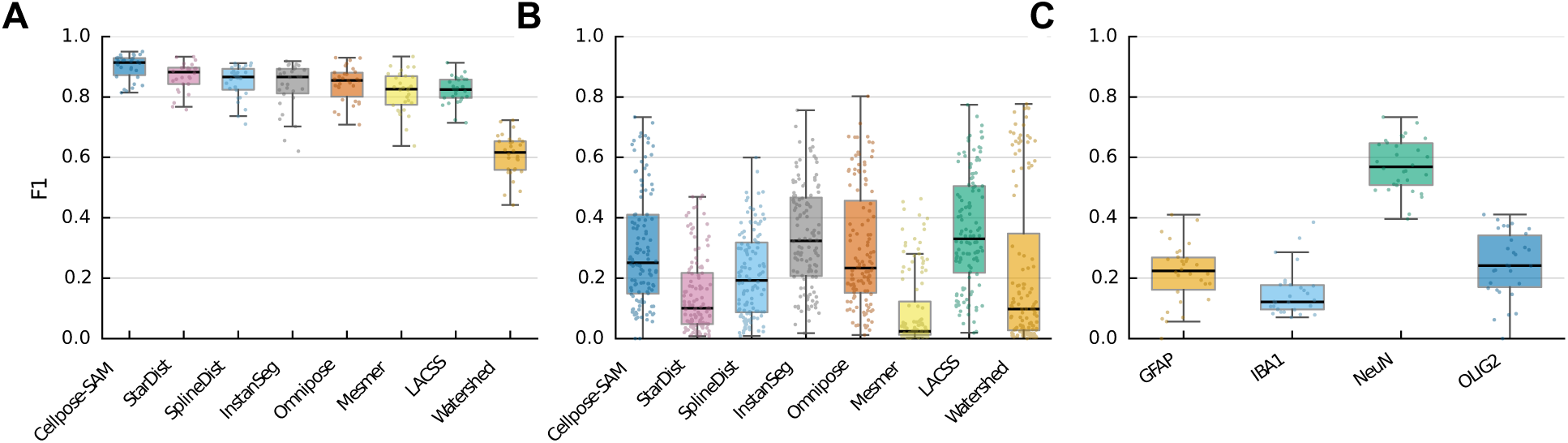
Benchmarking nuclear and cell segmentation with expert annotations. (A) Nuclear segmentation F1 for each benchmarked method. Each box summarizes F1 values computed per image, and each point is one image; colors identify methods. (B) Cell segmentation F1 for each benchmarked method, shown as in (A). (C) Cell segmentation F1 of zero-shot Cellpose-SAM for each marker; each box summarizes per-image F1 values, each point is one image, and colors identify markers.

All methods produced low F1 values for cell segmentation (Fig. 2B), and strong nuclear performance on DAPI did not transfer to the marker channel. For Cellpose-SAM, performance also varied by marker, with the highest F1 for NeuN and the lowest for IBA1 (0.57 versus 0.15; Fig. 2C). This difference is consistent with the marker-labeled structures: NeuN labels nuclei and perinuclear cytoplasm, with little labeling of neuronal processes, whereas IBA1 labels microglial perinuclear cytoplasm and radiating fine processes, which produces less regular cell-body boundaries.

Cellpose-SAM achieved the strongest nuclear F1 while maintaining a competitive cellular F1 among the benchmarked methods (Figs. 2A,B). Cellpose-SAM leverages the SAM vision transformer encoder to directly predict Cellpose vector flow fields (*25*) and supports efficient fine-tuning on small annotated datasets. We therefore selected Cellpose-SAM for fine-tuning of cell segmentation.

### Fine-tuning brain cell segmentation with expert annotations

We assigned each donor, with all of that donor’s annotated images, to the training, validation, or test set, so that no donor contributed images to more than one set and test performance reflected generalization to new donors. On the test set, zero-shot Cellpose-SAM achieved an F1 of 0.37. Marker-only fine-tuning increased F1 to 0.81, and adding the DAPI channel as a second input further increased F1 to 0.84 (Fig. 3A). Performance was stable across random seeds: five independent runs with identical hyperparameters produced F1 values between 0.83 and 0.84. For comparison, StarDist, a non-foundation model, reached 0.68 after fine-tuning on the same image split (Fig. 3A). Gains were consistent across markers, although performance was higher for neuronal and oligodendrocyte targets, likely because these cells were labeled by proteins primarily localized to the cell body, unlike astrocytes and microglia, which have complex processes labeled by GFAP and IBA1, respectively (Fig. 3B). In representative images, zero-shot Cellpose-SAM often missed or fragmented marker-positive cell bodies, whereas the fine-tuned model recovered them across markers (Fig. 4).

**Figure 3:**
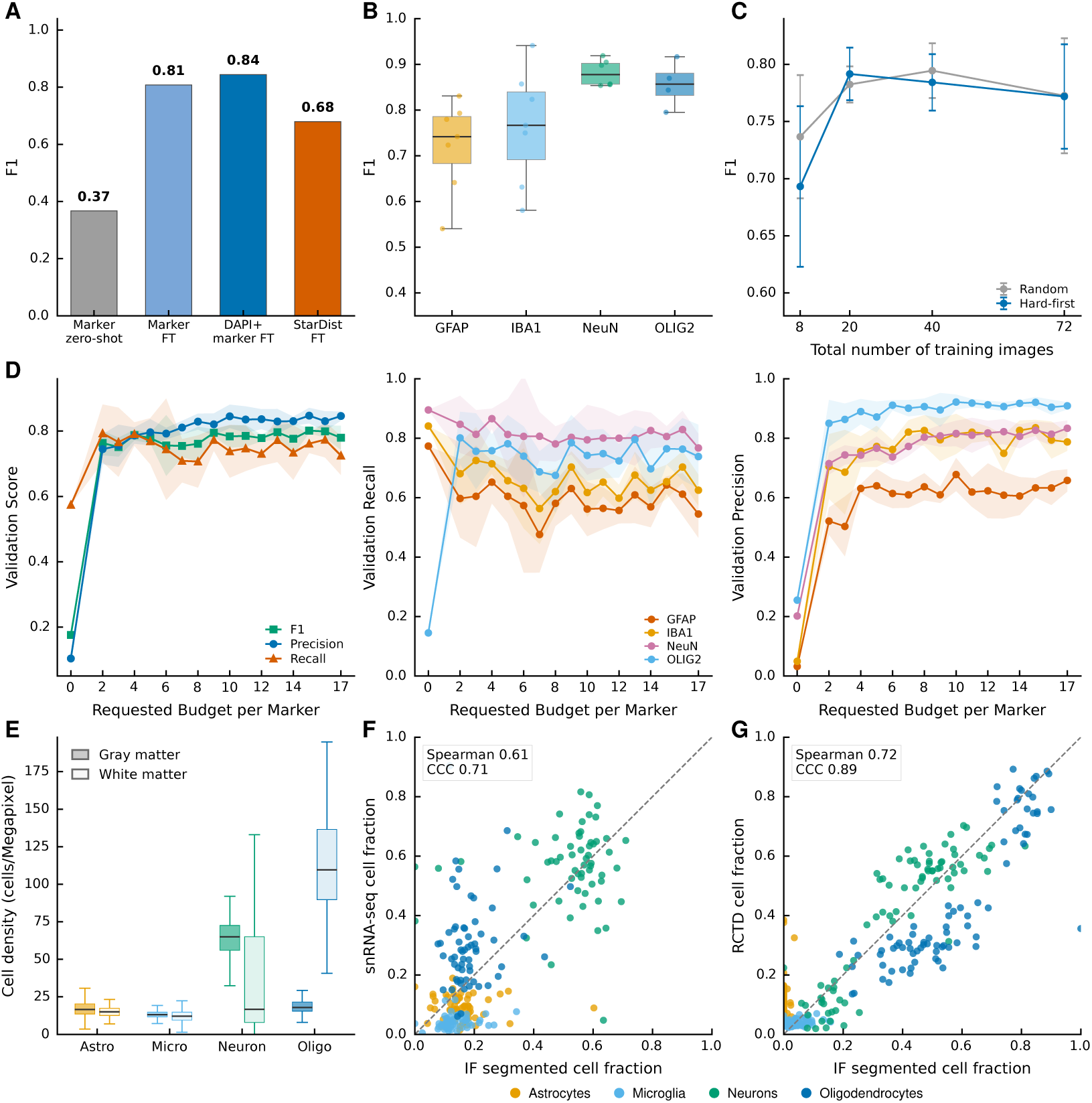
Performance of fine-tuned Cellpose-SAM for cell segmentation. (A) Test-set F1 for zero-shot, marker-only fine-tuned, DAPI + marker fine-tuned Cellpose-SAM, and fine-tuned StarDist. (B) Per-image F1 distributions across markers for fine-tuned Cellpose-SAM. (C) Active-sampling comparison showing validation F1 versus the total number of training images across four markers for random and hard-first image selection (mean ± SD of five repeats). (D) Validation performance versus the number of annotated training images per marker. (E) Marker-positive cell density estimated from fine-tuned segmentation across gray and white matter. White matter images were segmented using the model fine-tuned on gray matter annotations, since no white matter images were annotated. (F) Comparison of IF-derived and snRNA-seq cell fractions in gray matter. Each point is one donor and cell type. (G) Regional agreement between IF-derived and RCTD-deconvolved cell fractions in a Visium section (*9*). Each point is one 600-*μ*m bin and cell type. CCC: Lin’s concordance correlation coefficient.

**Figure 4:**
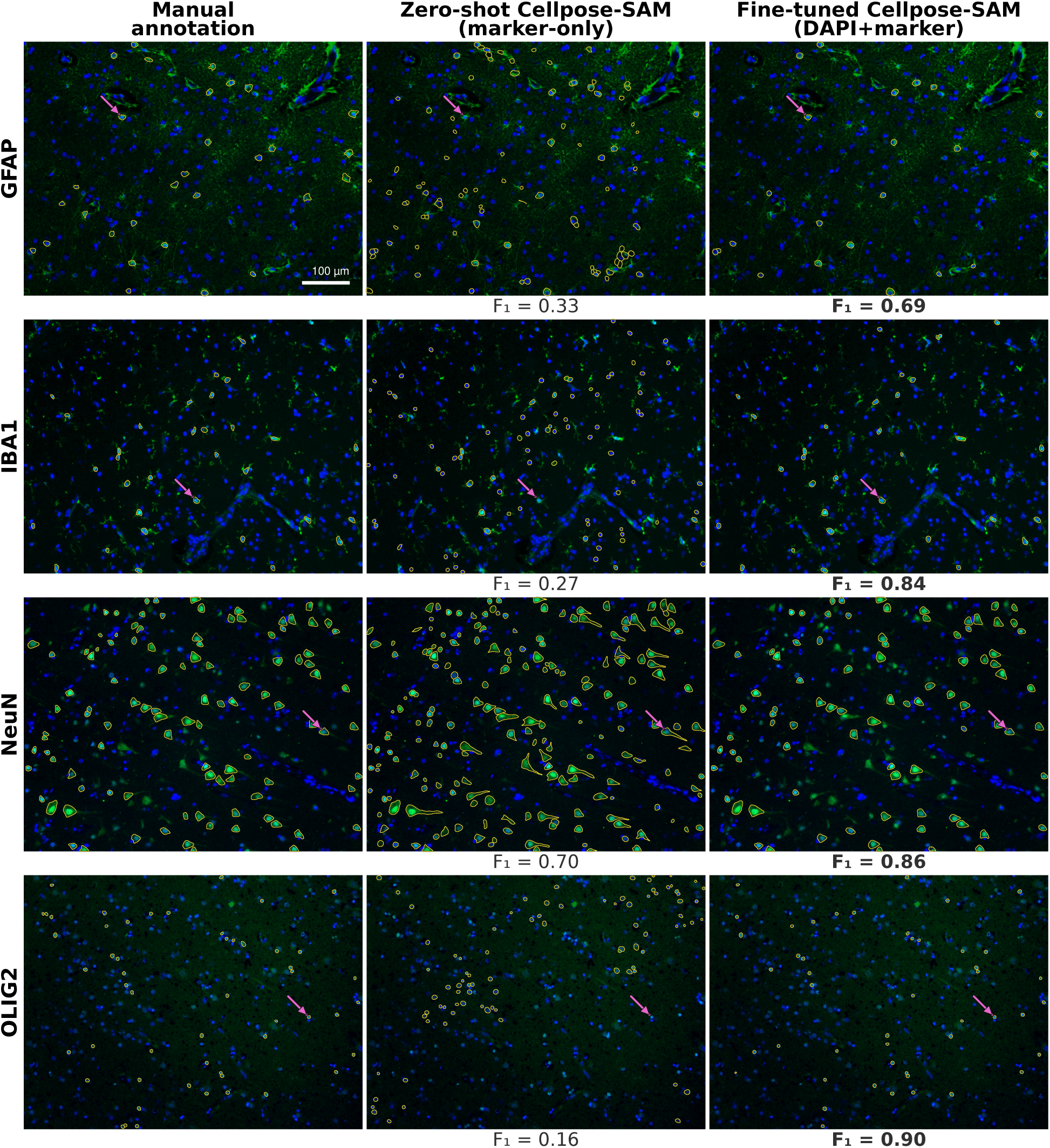
Cell segmentation in representative IF images for each marker. Columns show expert manual annotation (left), zero-shot Cellpose-SAM (middle), and fine-tuned Cellpose-SAM using the DAPI + marker input (right), with the per-image F1 score below each panel. Outlines denote annotated or predicted cell-body boundaries. Arrows mark the same cell in each row: cells missed by zero-shot segmentation in GFAP, IBA1, and OLIG2, and improved boundary delineation in NeuN. DAPI, blue; cell marker, green. Scale bar, 100 *μ*m; applies to all panels.

To determine how many annotated images were needed for fine-tuning, we evaluated model performance across progressively smaller training subsets. Prioritizing images with unstable zero-shot predictions for annotation (active sampling) did not improve F1 over donor-stratified random selection at matched annotation counts (Fig. 3C), suggesting that random sampling already covered the range of image difficulty at this scale. Few images were needed: with two annotated images per marker, the mean validation F1 reached 0.76 and approached 0.80 at larger budgets (Fig. 3D). A small annotation budget captured most of the observed performance gain.

### Cell compositions validated across platforms

We then evaluated whether the fine-tuned masks produced biologically plausible cell compositions, using three complementary checks: 1) tissue-compartment anatomy, 2) snRNA-seq, and 3) Visium spatial transcriptomics. First, we applied SPInfer to ROSMAP IF images from gray and white matter (*8*) and compared cell densities between the two compartments (Fig. 3E). Because all annotations were derived from gray matter, this comparison also tests whether the fine-tuned model generalizes to white matter. As expected, neuronal density was greater in gray matter, and oligodendrocyte density was greater in white matter. These cell density differences provided an internal anatomical check of the fine-tuned masks before downstream analyses.

Second, we compared the relative cell type composition of gray matter, predicted by fine-tuned Cellpose-SAM across all images from each donor, with snRNA-seq data from the same 60 donors in the ROSMAP-Compass atlas (*29*). Cell types were categorized as astrocytes (GFAP), microglia (IBA1), neurons (NeuN), or oligodendrocytes (OLIG2). The Spearman correlation between the two platforms’ cell fractions was 0.61 (Fig. 3F). In contrast, zero-shot masks were negatively correlated with snRNA-seq fractions (*ρ* = −0.54), and an existing watershed-based pipeline (*8*) showed weaker agreement (*ρ* = 0.55). Donor-aligned composition profiles across platforms are shown in Supplementary Fig. S1.

Third, we applied the same fine-tuned Cellpose-SAM model without retraining to IF images from an independent cohort in which Visium spatial transcriptomics was acquired on the same sections (*9*). Each section contained DAPI and four marker channels: astrocytes (GFAP), neurons (NeuN), oligodendrocytes (OLIG2), and microglia (TMEM119). The shared fine-tuned check-point, trained on all expert annotations, segmented each DAPI–marker pair separately, including TMEM119, which was absent from the training markers. We compared IF-derived composition with RCTD-deconvolved transcriptomic composition (*30*). Because spot-level composition estimates were sparse, we aggregated spots into a regular grid of 600-*μ*m bins (six Visium spot pitches; spot centers are 100 *μ*m apart). In section A1, one of the four Visium sections (see Materials and Methods), IF-derived and RCTD-deconvolved cell fractions showed regional-scale agreement (Spearman *ρ* = 0.72; Fig. 3G). Agreement in the remaining sections is shown in Supplementary Fig. S2 (B1: *ρ* = 0.48; C1: *ρ* = 0.62; D1: *ρ* = 0.53). In contrast, zero-shot masks showed little to no agreement in any section (*ρ*: A1, −0.04; B1, −0.13; C1, 0.02; D1, −0.16).

Together, these checks support the anatomical plausibility of the fine-tuned compositions and their agreement with transcriptomic data at the level of cell types and tissue regions.

### Pathology associations identified by fine-tuned segmenter only

We then applied SPInfer to 14,765 images from 69 ROSMAP donors spanning Braak stages 0– VI (*8*), segmenting 8.4 million nuclei. Of these, 557,301 were assigned to marker-positive cell bodies. To test whether relative cell fraction shifted with AD pathology, we fitted beta-binomial mixed-effects models to image-level cell counts. The response was the number of marker-positive cells out of all nuclei in an image, and each clinical variable was entered separately as a fixed effect. The beta-binomial distribution accommodated overdispersion, and a donor-level random intercept accounted for repeated images from the same donor (see Materials and Methods). Identical mixed-effects models were fitted to zero-shot and fine-tuned Cellpose-SAM masks.

Zero-shot masks produced two nominally significant associations, but neither survived Benjamini– Hochberg correction (*q* ≥ 0.23; Fig. 5A). In contrast, fine-tuned masks yielded four relative cell fraction associations at FDR (false discovery rate) < 0.05 (Fig. 5B). Three involved OLIG2: gray matter OLIG2 with Braak stage and CERAD (Consortium to Establish a Registry for Alzheimer’s Disease) score, and white matter OLIG2 with Braak stage. Because lower CERAD scores indicate more neuritic plaques, all three OLIG2 associations correspond to greater OLIG2-positive cell fractions with more severe AD pathology.

**Figure 5:**
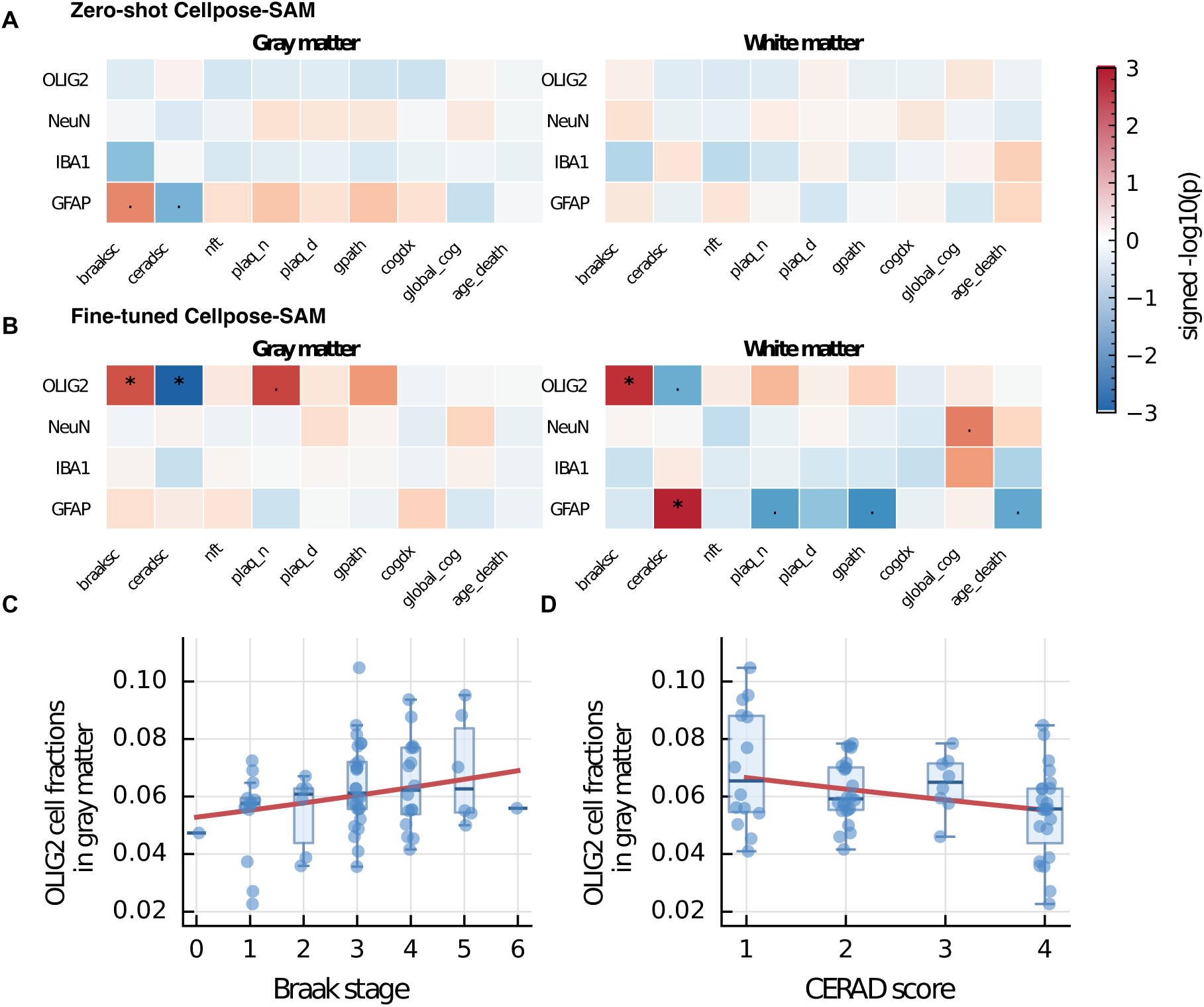
Associations between segmented cell fractions and AD variables. (A) No associations met the FDR threshold in the zero-shot Cellpose-SAM screen. (B) Four associations met the FDR threshold in the fine-tuned screen. ∗: FDR < 0.05; ·: *p* < 0.05. Color indicates − log_10_( *p*) from image-level beta-binomial mixed models, and the sign indicates effect direction, separated by gray and white matter. (C–D) Donor-level gray matter OLIG2-positive cell fraction across Braak stage and CERAD score; red lines show robust logit-linear fits for trend visualization. braaksc: Braak stage, a semiquantitative measure of neurofibrillary tangle (NFT) pathology; ceradsc: CERAD score based on neuritic plaques, coded in reverse with 1 indicating AD; plaq n: neuritic plaque burden; plaq d: diffuse plaque burden; gpath: global AD pathology burden, average of plaq d, plaq n, and nft; cogdx: cognitive diagnosis at death; global cog: global cognitive function.

OLIG2 is a nuclear transcription factor expressed in oligodendrocytes and their progenitors (*15, 16, 31*), so we interpret these associations as differences in the relative fraction of nuclei assigned to OLIG2-positive cells. The signal recurred across both pathology axes (Braak stage for neurofibrillary tangles and CERAD score for neuritic plaques) and across tissue compartments (gray and white matter). Donor-level visualizations of gray matter OLIG2 cell fractions showed corresponding trends across Braak stage and CERAD score (Figs. 5C,D). Analyses that directly linked cell-body morphology and spatial organization to AD clinical variables showed nominal associations, but none survived FDR correction (Figs. S3 and S4).

### Spatial graphs improve cell typing from nuclear features

Because single-marker IF labels only one cell type per image, we asked whether the remaining unlabeled nuclei in an image could be typed from DAPI nuclear features and local spatial context alone. We represented each image as a Delaunay graph (*32*) over all DAPI-positive nuclei (Supplementary Fig. S5). Nuclei assigned to GFAP-, IBA1-, NeuN-, or OLIG2-positive cell bodies carried one of four cell-type labels, while the remaining DAPI-positive nuclei were kept as unlabeled context nodes during message passing. Valid Delaunay graphs were constructed for the 13,280 images with more than three marker-positive nuclei.

Each nucleus was described by 37 CellProfiler features (*33*) computed on its nuclear mask: 24 morphological and 13 DAPI-intensity measurements (Fig. 6A; see Materials and Methods). The classifier received no marker-channel information. Cell typing from DAPI-derived features and spatial context offers a route to cell-type mapping when multiplexed marker panels are unavailable. Using these nuclear features alone, a multilayer perceptron (MLP) and an XGBoost model achieved test-set macro-F1 scores of 0.70 and 0.68, respectively. Adding Delaunay nuclear graphs with a two-layer GATv2 (graph attention network) encoder (*34*) raised macro-F1 to 0.89, averaged over five random seeds using donor-disjoint data partitions (Fig. 6B). The node features were held constant between models, showing that the graph-based model classified cells more effectively than models using per-nucleus features alone.

**Figure 6:**
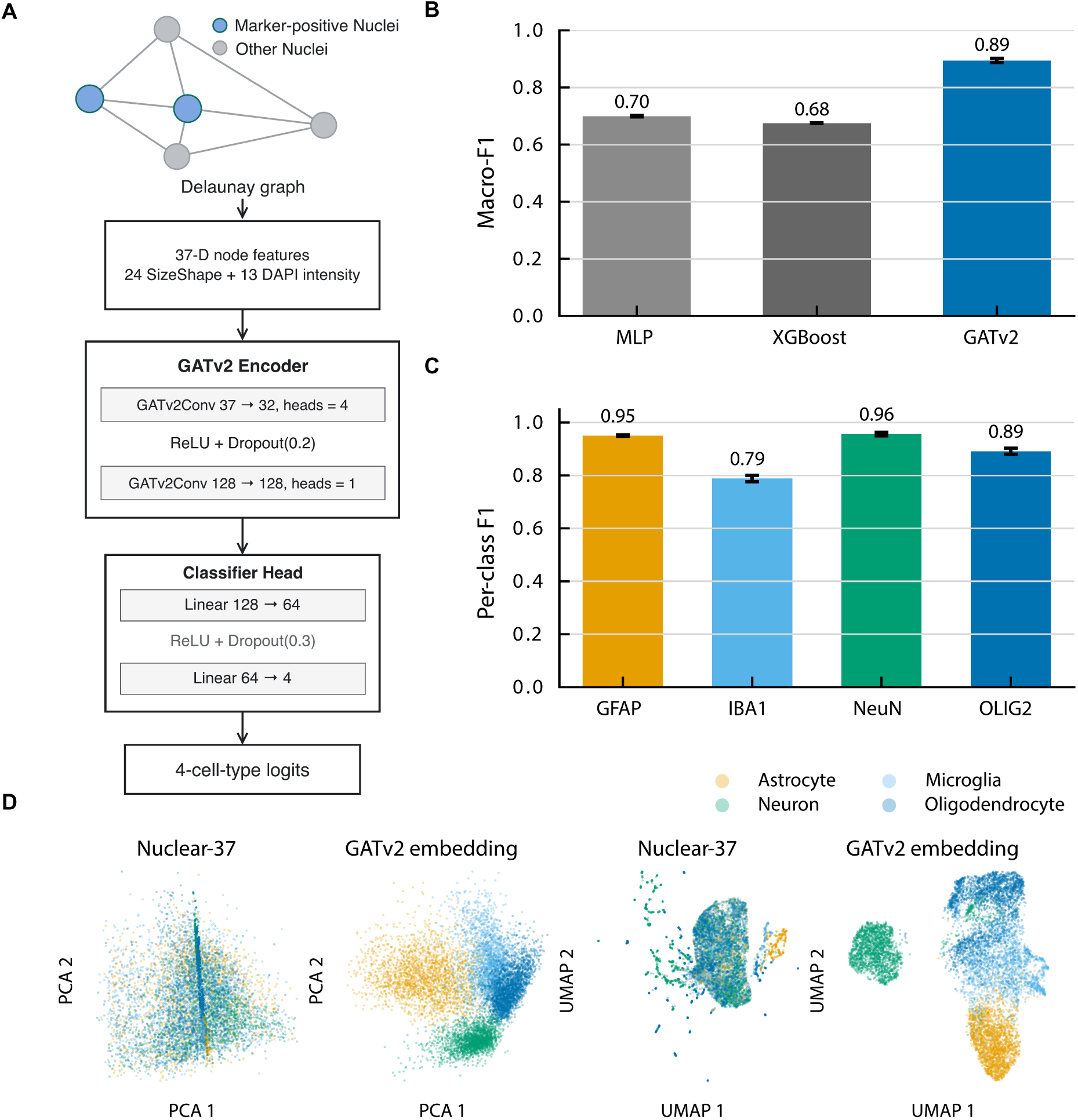
Fine-tuned segmentation enables spatial graph construction and graph-based cell typing. (A) GATv2 applied to Delaunay graphs of nuclei for cell type classification. (B) Four-class macro-F1 for non-graph baselines and GATv2 models using the same 37 CellProfiler features per nucleus. (C) Per-class GATv2 F1 across random seeds. (D) PCA and UMAP visualization of nuclear features and GATv2 embeddings on the test set, colored by cell type.

Per-class F1 was highest for NeuN-positive neurons (0.96) and GFAP-positive astrocytes (0.95), followed by OLIG2-positive oligodendrocytes (0.89) and IBA1-positive microglia (0.78) (Fig. 6C). In principal component analysis (PCA) and uniform manifold approximation and projection (UMAP) plots, cell types separated more clearly in the GATv2 embedding than in the raw 37-feature space (Fig. 6D). With two GATv2 layers, each prediction drew on a nucleus’s direct Delaunay neighbors and their neighbors (Fig. 6A). The gain from this two-hop context is consistent with the biological expectation that cell identity is partly reflected in the composition and arrangement of the surrounding tissue.

### Linking segmented cell morphology to gene expression

Using the fine-tuned cell-body masks, we asked whether donor-level morphology covaried with gene expression in the corresponding cell type. For each marker class in gray matter, we computed the same 37 CellProfiler features on marker-positive cell-body masks, summarized them as donor medians, and related them to pseudobulk snRNA-seq profiles of the same cell type (raw counts summed across nuclei of that cell type within each donor) from 53 to 57 donors per class, adjusting for age, postmortem interval, and technical covariates (see Materials and Methods).

At *q* < 0.05, corrected across all gene–feature–class tests, we identified 2,267 associations involving 602 genes. More than half involved IBA1-positive cells (1,310), followed by GFAP-positive (470), NeuN-positive (357), and OLIG2-positive (130) cells (Fig. 7A).

**Figure 7:**
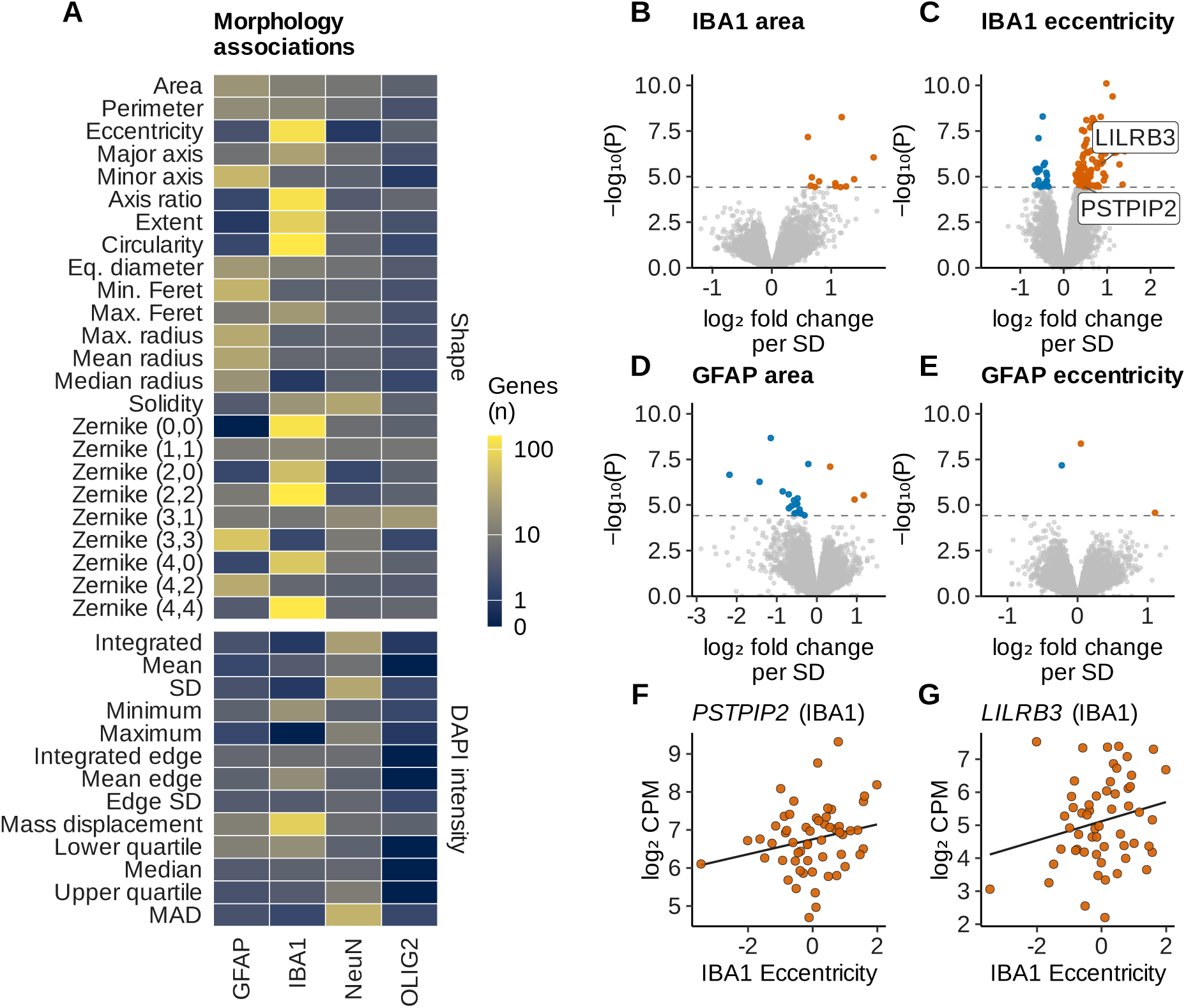
Donor-level cell-body morphology covaries with gene expression in gray matter. (A) Number of genes associated with each of the 37 features in each cell type, colored on a log(1 + *n*) scale. FDR was controlled across all gene–feature–class tests. (B–E) Volcano plots for cell-body area and eccentricity in IBA1-positive (B, C) and GFAP-positive (D, E) cells. Each point is one gene. The *x*-axis shows the log_2_ fold change per SD increase in the donor-median feature, and the *y*-axis shows − log_10_( *p*). Orange and blue points indicate positive and negative associations with global FDR < 0.05, gray points are not significant, and dashed lines mark the corresponding *p*-value threshold. (F, G) IBA1 eccentricity versus *PSTPIP2* and *LILRB3* expression. Both genes, labeled in (C), were selected after the screen for display based on prior experimental evidence linking them to cell shape. Lines are least-squares fits.

Within IBA1-positive cells, the six features associated with the largest number of genes were Zernike moment (2,2) (142 genes), circularity (136), Zernike (4,4) (135), Zernike (0,0) (124), axis ratio (121), and eccentricity (116) (Fig. 7A). These features were strongly correlated across donors, and each correlated with eccentricity at |*ρ*| ≥ 0.84 (Fig. S6), indicating that they largely capture a shared morphological axis of cell elongation. Eccentricity and axis ratio, for example, shared 105 associated genes. Area was only weakly to moderately correlated with these features (|*ρ*| ≤ 0.67) but strongly correlated with other size measures.

We therefore focused on two interpretable representatives, area for cell size and eccentricity for elongation, in the two cell types with the most associations. In IBA1-positive cells, eccentricity was associated with 116 genes, whereas area was associated with only 12 (Figs. 7B,C). GFAP-positive cells showed the reverse, with more associations for area (21) than for eccentricity (3; Figs. 7D,E). Two of the eccentricity-associated genes have been linked experimentally to cell shape in macrophages and monocytes. In mouse macrophages, increased *PSTPIP2/MAYP* expression was associated with increased cell spreading and elongation (*35*). In primary human CD14-positive monocytes, agonistic *LILRB3* activation produced a more elongated morphology (*36*). Across ROSMAP donors, those with more elongated IBA1-positive cells showed higher *PSTPIP2* expression (*q* = 0.048; Fig. 7F) and higher *LILRB3* expression (*q* = 0.007; Fig. 7G).

## Discussion

SPInfer demonstrates the value of fine-tuning for cell segmentation and downstream pathology analysis in brain tissue. Several methods performed well on nuclear segmentation, but zero-shot segmentation of marker-positive cell bodies remained a major challenge. Fine-tuning Cellpose-SAM more than doubled test-set cell segmentation F1. Using the same pathology models, the fine-tuned pipeline identified four FDR-significant associations, whereas the zero-shot pipeline identified none. These results establish segmentation as an important component of cohort-scale pathology inference and connect annotation-based model evaluation with downstream biological analysis.

Several analyses support the biological validity of the fine-tuned masks. Segmentation-derived cell densities recovered expected gray–white matter differences. IF-derived compositions showed concordance with snRNA-seq data from the same donors and regional concordance with independently acquired Visium spatial transcriptomics data. In contrast, zero-shot masks showed no or negative concordance with either platform, indicating that this agreement depended on segmentation accuracy. Agreement in cell-type composition and regional variation supports the use of SPInfer-derived fractions for cohort-scale spatial proteomic analysis.

The pathology association study highlighted OLIG2-positive fractions as a recurring correlate of AD pathology. This convergence across pathology axes and tissue compartments is consistent with recent work implicating oligodendrocyte involvement in AD progression (*16*). Our protein-level observation provides an independent line of evidence at single-cell image resolution. SPInfer also extends naturally from cell counting to spatial analysis. Using the same per-nucleus features, a graph attention model over Delaunay nuclear graphs improved the macro-F1 compared with non-graph models. This finding is biologically plausible: different cell types are embedded in characteristic tissue microenvironments, and two-hop message passing through the Delaunay graph captures these spatial regularities. The donor-level morphology analysis identified coordinated variation between cell-body features and RNA expression, with the largest number of associations observed for IBA1-positive cells.

This study has several limitations that motivate future work. The associations with AD pathology are observational and do not establish causality or temporal sequence. Similarly, the snRNA-seq comparison shows agreement across cell types but does not establish donor-specific agreement. The morphology–expression analysis has a related constraint: imaging and RNA came from the same donors and cell types but not from the same cells. Subtype composition, tissue handling, and measurement variation could therefore contribute to the observed associations. Because the most frequently associated shape features were strongly correlated, the number of significant gene–feature associations may not correspond directly to the number of independent biological processes. In the graph-based cell-typing analysis, the single-marker design couples image identity with marker identity. We centered DAPI intensity features within each image to account for marker-associated shifts, although residual image-level effects cannot be entirely excluded.

In future studies, replication in independent AD cohorts will be needed to test whether the OLIG2 cell-fraction associations and the morphology–expression signal generalize beyond ROSMAP. Determining whether the IBA1 morphology–expression associations reflect a cell-autonomous mechanism will also require experiments in human brain myeloid cells. For the graph-based cell typing, multiplexed labels on matched nuclei would provide a stronger test of the single-marker design and separate biological neighborhood structure from image-level acquisition effects. SPInfer itself currently segments two-channel 2D images, and both the number of channels and the dimensionality could be extended. With multichannel inputs, such as high-plex IF panels, it could segment multiple cell types jointly and measure marker co-expression directly.

SPInfer provides a practical workflow for cohort-scale spatial proteomics. A small number of annotated images per marker captures most of the attainable performance gain, lowering the barrier to adoption in new cohorts or marker panels (*37*). To support reproducibility, we release our expert annotations as a valuable reference for benchmarking and fine-tuning brain cell segmentation models, alongside model weights, environment specifications, and reproduction workflows. In human brain tissue, SPInfer improves cell-body segmentation and links the resulting measurements to pathology, graph-based cell typing, and donor-level gene expression. The workflow connects annotation, model adaptation, cohort deployment, and hierarchical statistical analysis, providing a basis for applications in other tissues and disease conditions (*5, 6, 9, 38*).

## Materials and Methods

### Datasets and preprocessing

#### Human brain IF images and annotations

Brain tissue samples were obtained from the Religious Orders Study and Rush Memory and Aging Project (ROSMAP) (*7*). IF images of the dorsolateral prefrontal cortex were generated as previously described (*8*); sections were stained with DAPI and labeled for one of five markers (GFAP, IBA1, NeuN, OLIG2, or PECAM) and imaged at 20× on a Zeiss Axio microscope. Full-cohort inference covered 14,765 images (1,040 × 1,388 pixels each) from 69 donors spanning Braak stages 0–VI, with gray- and white-matter labels available for each image.

Expert annotations were generated by annotators with experience in human brain neuroanatomy, histopathology, and IF imaging. Digital micrographs were analyzed in ImageJ, and annotators were blind to all subject characteristics. Micrograph brightness and contrast were altered to improve object identification and annotation. For nuclear segmentation, annotators manually traced boundaries visible in the DAPI channel. For cell segmentation, marker-positive cells were defined as those with immunofluorescence signal consistent with established morphological features and a visible DAPI-stained nucleus; annotators then traced the corresponding cell body. PECAM-labeled images were excluded from the cell segmentation benchmark and the downstream tasks because PECAM labels continuous tubular vasculature rather than discrete cell bodies.

For nuclear segmentation, 30 DAPI images containing 14,678 nuclei were annotated. For cell segmentation, we annotated 120 cell-marker images (30 per marker), containing 5,832 marker-positive cell bodies: 1,487 OLIG2, 2,771 NeuN, 836 IBA1, and 738 GFAP. All annotated images were from gray matter; no white matter annotations were generated.

#### Image preprocessing

ROSMAP IF images were converted into standardized inputs by maximum-intensity projection of z-stacks, channel-wise intensity clipping to the 0–98th percentile, and rescaling to [0, 1]. When a method explicitly required additional normalization, we followed the recommended default workflow. No method-specific parameter tuning or bespoke preprocessing was introduced beyond these default or mandatory steps. DAPI + marker two-channel inputs for Cellpose-SAM consisted of the DAPI channel and the corresponding marker channel; single-channel inputs used only the marker channel.

#### Single-nucleus RNA-seq data

For cross-platform gray matter benchmarking, we used donor-level single-nucleus gray matter cell fractions processed from the published ROSMAP-Compass snRNA-seq atlas (*29*) and restricted analysis to donors shared with the IF cohort. For each donor, we computed gray matter cell-type fractions from the fine-tuned segmentation masks by pooling marker-positive cell counts across all gray matter images and dividing each cell-type count by the total across the four cell types, ensuring that the four fractions summed to one. For comparison, the same fractions were also computed from zero-shot Cellpose-SAM masks and from an existing watershed-based pipeline with marker-intensity filtering. Cell types were merged into the four shared categories: astrocytes, microglia, neurons, and oligodendrocytes. Agreement between platforms was summarized using Spearman correlation and Lin’s concordance correlation coefficient.

#### Spatial transcriptomics data

Four dorsolateral prefrontal cortex sections (V10B01-087 A1, B1, C1, D1) were processed on the 10x Genomics Visium platform with paired IF imaging (*9*). Each IF image contained a DAPI channel together with four marker channels (GFAP, NeuN, OLIG2, and TMEM119). Each section was cropped to the bounding box of its Visium spot centers with a fixed 600-pixel external margin; tissue farther outside this region was excluded before scaling and inference. The fine-tuned Cellpose-SAM model was applied to each DAPI–marker pair without additional retraining, using Cellpose default normalization, a flow threshold of 0.4, and a cell-probability threshold of 0.0. Segmented centroids were mapped to Visium spots after registration of the IF and Visium coordinate systems, using centroid-in-circle assignment at a spot radius of 65.5 px (*9*), equal to half the 131-pixel full-resolution spot diameter. Image-derived composition was compared with reference RCTD deconvolution (*30*) after binning on a regular grid with a 600-*μ*m bin width (six times the 100-*μ*m Visium center-to-center spot distance, converted to pixels for each section using its median nearest-neighbor spot distance); bins containing fewer than eight spots were excluded. Zero-shot Cellpose-SAM was evaluated with identical preprocessing and binning for comparison.

#### Benchmarked segmentation methods

We evaluated eight publicly released segmentation tools spanning classical image processing, task-specific convolutional networks, and foundation models. All methods were executed using their original Python packages or APIs, with their recommended pretrained weights; no additional training or fine-tuning was performed during benchmarking, unless indicated as fine-tuned.

#### Watershed

The watershed transform interprets an intensity or distance map as a topographic surface and iteratively floods local minima until catchment basins converge, thereby partitioning the image into instances. For nuclei, seeds were generated from local maxima of a Euclidean distance transform of Otsu-thresholded DAPI; for cell bodies, seeds were derived from Sobel gradients of the corresponding marker channel. We used the skimage.segmentation.watershed API with connectivity = 1 and default compactness, relying solely on the gradient magnitude image as the flooding surface. Boundary fragments were closed with a single morphological dilation–erosion cycle using a disk structuring element of radius 3 px to create contiguous masks.

#### StarDist

StarDist models each object as a star-convex polygon (*17*) parameterized by a fixed number of radial distances emanating from every pixel. A U-Net backbone jointly predicts object probability and 32 radial distances, allowing simultaneous detection and shape reconstruction. We employed the “2D versatile fluo” checkpoint via the stardist API with default non-maximum suppression (IoU threshold 0.3) and Gaussian-based probability thresholding.

#### SplineDist

SplineDist extends StarDist by representing contours as cubic B-splines defined by learnable control points, thereby removing the star-convexity constraint (*18*). We used the publicly released pretrained weights, which provide models trained with different numbers of control points. The checkpoint used in our code was loaded from the extracted splinedist models bundle and executed through the TensorFlow API; architectural details (including the number of control points) followed the checkpoint defaults. Predicted spline curves were rasterized; overlapping instances were resolved by retaining pixels with the highest object probability. No additional parameter tuning was performed.

#### Omnipose

Omnipose formulates segmentation as vector-flow regression, directing each pixel toward its cell’s medial axis, enabling robust delineation of irregular or branched morphologies. We used the cellpose API with MODEL TYPE = "auto" and omni=True, which selects cyto according to the library’s default logic, with a flow threshold of 0.4 and default diameter estimation. Objects smaller than 15 px (nuclei) or 30 px (cells) were discarded, and all other hyperparameters were left at their default settings.

#### LACSS

The location-aware cell segmentation system (LACSS) combines a Mask R-CNN backbone with auxiliary location-consistency losses to leverage weak supervision (*22*). We used the TissueNet-pretrained checkpoint available in the lacss package and invoked the predict API with a default confidence threshold of 0.5. No user-defined seeds or retraining were introduced.

#### Mesmer

Mesmer is a dual-decoder network designed for multiplexed tissue imaging that outputs nuclear and cell masks concurrently (*20*). The DeepCell 0.12 pretrained weights were executed through the DeepCell-tf API at the recommended physical resolution of 0.5 *μ*m/px. Histogram matching to the TissueNet intensity profile was automatically applied by the API; subsequent instance masks were accepted as-is, except for removing objects outside the 50–5,000 px^2^ range.

#### InstanSeg

InstanSeg is an embedding-based instance segmentation pipeline that predicts a seed map alongside positional and conditional embeddings, then assigns pixels to instances by comparing per-pixel embeddings to optimally selected seed embeddings (*39*). We ran the pre-trained fluorescence checkpoint (fluorescence nuclei and cells) via the instanseg-torch InstanSeg class using the default eval small image inference pathway.

#### Cellpose-SAM

Cellpose-SAM leverages the SAM vision transformer encoder to directly predict Cellpose vector flow fields, supporting zero-shot segmentation (*25*). In our implementation, we ran the public API using the default inference settings and enabled automatic diameter estimation (i.e., no fixed diameter and no explicit tiling configuration in the call). No retraining or hyperparameter tuning was performed for this benchmark.

### Fine-tuning protocol

#### Training objective and channel configuration

Fine-tuning retained the Cellpose training objective without modification; only the input tensor and the supervision target were specific to this study. Let *X* ∈ R*^C^*^×^*^H^*^×^*^W^* denote an input image over the pixel domain Ω, with *C* = 2 for the DAPI + marker configuration and *C* = 1 for the marker-only configuration. The network *f_θ_* predicts a two-component spatial flow field *V*^^^ ∈ R^2×^*^H^*^×^*^W^* and a logit-transformed probability that each pixel belongs to any cell *p*^ ∈ R*^H^*^×^*^W^* . Targets were derived from the expert annotations: the reference flow *V*^∗^ was obtained by the simulated-diffusion procedure of Cellpose applied to the annotated instance masks (*19*), and *m*^∗^ = **1**{*u* lies within an annotated marker-positive cell body}. We minimized

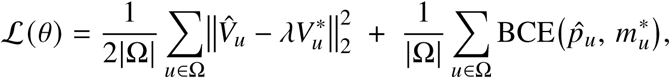

where BCE is binary cross-entropy with logits and *λ* = 5 is the default target-flow scaling of the Cellpose implementation.

The two configurations differ only in *C*: input channels are stacked along the channel dimension and consumed at the patch-embedding layer of the SAM encoder, while all subsequent parameters, the flow and probability heads, and L are identical. Critically, the DAPI channel appears only in *X* and never in *V*^∗^ or *m*^∗^: supervision is defined entirely by marker-delineated cell-body boundaries, so the two channels play asymmetric roles. The marker channel determines which instances exist and where their boundaries lie, whereas the DAPI channel supplies structural evidence of nucleus location that constrains instance separation where the marker signal is weak or where adjacent cell bodies abut.

#### Architecture and training

The annotation set was split at the donor level into 72 training, 24 validation, and 24 test DAPI-marker pairs during fine-tuning. There was no donor overlap between partitions, no image-hash collisions, and no near-duplicate image pairs. Training was performed on the 72-image training partition with standard augmentations, including rotation, flipping, and scaling. Models were trained for 500 epochs with a learning rate of 10^−4^, a weight decay of 10^−5^, and a batch size of 8. The checkpoint with the best validation loss was selected for evaluation. For comparison, we also fine-tuned StarDist (*17*) on the same donor-stratified training partition. StarDist used a 32-ray two-dimensional configuration with a (2, 2) grid, 256 × 256 training patches, and initialization from the 2D versatile fluo pretrained model when channel dimensions matched. Because the StarDist configuration used here accepts a single input channel, the normalized DAPI and marker channels were fused into a single composite input channel by pixel-wise averaging before training. The model was trained for 80 epochs with a learning rate of 3 × 10^−4^, a batch size of 2, 72 steps per epoch, and the same validation partition used for threshold optimization before test-set evaluation.

#### Seed stability

To assess reproducibility, five independent training runs were conducted with all hyperparameters fixed. Each model was evaluated at a shared operating point, defined as the median of the validation-selected thresholds across runs, to isolate the effect of random initialization from threshold selection.

#### Label efficiency

To assess the relationship between annotation budget and performance, models were trained on stratified subsets of the training partition containing 2 to 17 images per marker. Each budget level was evaluated on the full validation partition using the same threshold-selection protocol.

#### Active sampling

To test whether annotation could be targeted more efficiently, we compared random image selection with a hard-first sampling rule. Candidate training images were first scored based on the stability of zero-shot Cellpose-SAM predictions. The same image was segmented at two operating points, flow threshold/cell-probability threshold (0.4, 0.0) and (0.2, −0.5), and difficulty was defined as the normalized change in predicted instance count, |*n*_1_−*n*_2_|/(*n*_1_+*n*_2_+1), where *n*_1_ and *n*_2_ denote the predicted cell counts in the two runs. Images with larger scores were considered harder because their predicted object counts were more sensitive to the inference threshold; images with smaller scores were considered easier. For each marker and annotation budget, random sampling uniformly selected the required number of images, whereas hard-first sampling selected from the highest-scoring pool within that marker class. Models from both strategies were trained under the same fine-tuning and validation protocol and compared by validation F1 at matched annotation counts.

### Downstream pathology analysis

#### Image-resolved mixed models

The primary association analysis used image-level observations rather than donor averages. After technical-replicate deduplication to one observation per image, we retained images with non-zero nuclear counts, valid marker-positive counts, and complete donor and clinical metadata. For each marker, tissue compartment (gray and white matter), and clinical variable, we fitted a separate beta-binomial mixed-effects model. Let *m_i_* denote the marker-positive count, *n_i_* the total nuclear count, and *x_i_* the clinical variable for image *i*. We modeled

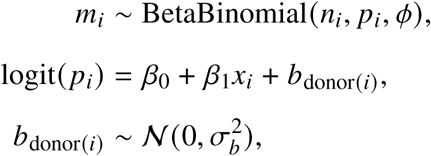

where *φ* denotes the overdispersion parameter and *b*_donor(_*_i_*_)_ is a donor-specific random intercept. This formulation preserves the image-specific nuclear denominator while accommodating overdispersion relative to a binomial model and within-donor clustering across multiple images. Clinical variables were entered one at a time (*40*).

### CellProfiler feature extraction

We computed a common set of 37 features with CellProfiler’s size-shape and intensity modules (*33*) and applied it to two mask sources from the fine-tuned Cellpose-SAM model. The features comprised 24 morphological measurements (area, perimeter, eccentricity, major axis length, minor axis length, solidity, extent, equivalent diameter area, axis ratio, circularity, maximum radius, mean radius, median radius, minimum Feret diameter, maximum Feret diameter, and nine Zernike moments z^m^_n_ with (*n*, *m*) ∈ {(0, 0), (1, 1), (2, 0), (2, 2), (3, 1), (3, 3), (4, 0), (4, 2), (4, 4)}) and 13 DAPI intensity measurements (integrated, mean, standard deviation, min/max, edge intensity statistics, mass displacement, quartiles, and median absolute deviation), each measured within the mask being described. Each Zernike moment z^m^_n_ describes the mask’s shape after mapping it onto a unit disk, where small *n* describes the overall outline (e.g., round versus oval) and larger *n* captures smaller bumps and indentations along the edge, and *m* counts how many times a pattern repeats around the cell outline (e.g., *m* = 2 corresponds to two opposite bulges, as in an oval; *m* = 4 to four lobes). We used the rotation-invariant magnitudes |z^m^_n_|; for example, |z^m^_n_| is near zero for a round cell and larger for an elongated, oval-shaped cell. For graph-based cell typing (Fig. 6), the features were computed on the nuclear mask of every DAPI-positive nucleus. For the morphology analyses (Fig. 7 and Figs. S3, S4, and S6), the same features were computed on marker-positive cell-body masks. Only DAPI-channel intensity was used; marker-channel intensity was excluded because single-marker IF couples the marker channel to the image’s target class, and including it would create direct label leakage.

### Graph neural cell-type classification

#### Graph construction

Graph neural analyses used the full-cohort Delaunay graphs derived from the fine-tuned masks. Each image was represented by a single graph whose nodes were all DAPI-positive nuclei, each with an assigned centroid. Delaunay triangulation was constructed after excluding nuclei within 32 pixels of the image border. Edges were treated as bidirectional spatial adjacencies. Marker-positive nuclei were assigned the cell-type label corresponding to the image marker, whereas non-target DAPI-positive nuclei were retained as unlabeled context nodes for message passing and excluded from the supervised loss.

#### Node features and batch correction

The main graph feature set contained 37 per-nucleus features: 24 CellProfiler size-shape measurements and 13 DAPI intensity measurements. Because single-marker IF imaging couples image identity with marker identity, we screened for marker-associated DAPI shifts at the image level. Raw DAPI intensity showed a marker effect, so the primary feature set applied within-image centering only to the 13 DAPI intensity dimensions. The 24 morphology dimensions were not image-centered.

#### Models and training

We employed a lightweight two-layer GATv2 network (*34*) with 51,845 trainable parameters. The first layer applied 4-head attention with 32 hidden units per head, yielding a 128-dimensional intermediate representation after concatenation. ReLU activation and feature dropout (0.2) were applied between layers, and the GATv2 layers used attention dropout of 0.2. The second layer used a single attention head and an output dimension of 128. Node embeddings were passed through a classification head comprising a linear layer (128 to 64), a ReLU activation, dropout (0.3), and a final linear layer (64 to 4), producing class logits. Cross-entropy loss was applied only to marker-positive nuclei; non-target DAPI-positive nuclei participated in message passing as context but did not contribute to the loss. Optimization used Adam with differential learning rates for the encoder and classification head, weight decay 10^−5^, batch size 32 tile graphs, 50 epochs, and early stopping on validation macro-F1. Non-graph baselines included a 37-feature MLP and XGBoost (*41*). All models used donor-disjoint train/validation/test partitions, and the GATv2 result was summarized across five random seeds.

### Evaluation metrics

We evaluated segmentation using object-matching metrics, pathology associations using mixed-effects models, cross-platform agreement using correlation and concordance measures, and cell-type classification using class-specific F1 and macro-F1.

#### Segmentation benchmarking and fine-tuning

Predicted instance masks were compared against expert annotations with the stardist.matching module (*17*), which performs one-to-one matching between annotated objects *G_i_* and predicted objects *P_j_* by descending overlap subject to an IoU threshold *τ*. For any candidate pair, overlap was defined by the Jaccard index

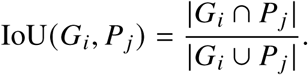

At a fixed threshold *τ* = 0.5, matched pairs were counted as true positives TP*_τ_*, unmatched predictions as false positives FP*_τ_*, and unmatched ground-truth objects as false negatives FN*_τ_*. We then computed

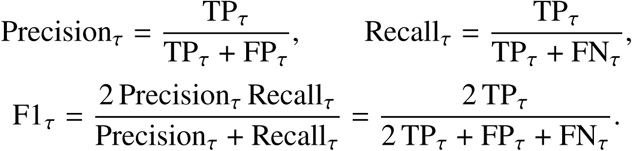

#### Graph-based cell-type classification

For the classification task, we report class-specific F1 scores,

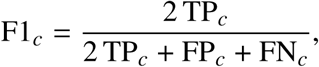

and macro-F1 across *C* = 4 classes,

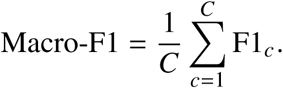

### Morphology-associated gene expression

Gene-level count models were fitted with edgeR (version 4.6.3) using a negative-binomial generalized linear model and robust quasi-likelihood inference (*42, 43*). Low-expression genes were removed separately for each model with filterByExpr, using the full design matrix and min.count=10. Library sizes were recomputed after filtering, and trimmed mean of M-values (TMM) normalization was applied (*44*). The retained gene counts were 19,070 for GFAP, 14,054 for IBA1, 26,075 for NeuN, and 21,319 for OLIG2 in the area/eccentricity models. One model was fitted for each of the 37 features and the 4 matched classes. The design was

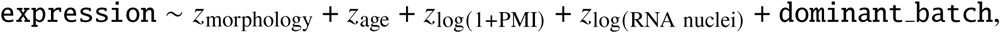

with the TMM-adjusted library size included as an offset on the log-count scale. Continuous predictors were centered and divided by their sample standard deviations among the included donors. Sequencing batch was a categorical covariate derived from the RNA metadata; dominant batch denoted the batch that contributed the most nuclei for a given donor. Sex was excluded because all included donors were male. No cell-composition or subtype-fraction covariates were added, and morphology features were tested separately rather than mutually adjusted. Every design matrix was checked for full rank and at least 10 residual degrees of freedom. Dispersion estimation and quasi-likelihood fitting used estimateDisp(…, robust=TRUE) and glmQLFit(…, robust=TRUE). The morphology coefficient was tested using glmQLFTest. Reported log_2_ fold changes quantify expression change per one-standard-deviation increase in the donor-median feature. A single Benjamini–Hochberg adjustment was applied to all 2,979,166 gene–feature–class *p* values from the 148 models (37 features × 4 classes); *q* < 0.05 defined significance. No additional within-model correction or fold-change cutoff was used.

## Acknowledgments

We thank the participants of the Religious Orders Study and Rush Memory and Aging Project (ROSMAP). ROSMAP is supported by P30AG10161, P30AG72975, R01AG15819, R01AG17917, U01AG46152, and U01AG61356. The results published here are in whole or in part based on data obtained from the AD Knowledge Portal (NIH-NIA, U24AG061340). snRNA-seq data were generated from postmortem brain tissue provided by the ROSMAP cohort at Rush Alzheimer’s Disease Center, Rush University Medical Center, Chicago, and were funded by NIH grants U01AG061356, RF1AG057473, U01AG046152, R01AG066831, and U01AG072572. This research was supported in part by the University of Pittsburgh Center for Research Computing and Data, RRID:SCR 022735, through the resources provided. Specifically, this work used the H2P cluster, which is supported by NSF award number OAC-2117681.

## Ethics approval and consent to participate

The Religious Orders Study and Rush Memory and Aging Project were approved by an Institutional Review Board of Rush University Medical Center. All participants signed an informed consent, a repository consent, and an Anatomic Gift Act. ROSMAP data were obtained through the Rush Alzheimer’s Disease Center Resource Sharing Hub under a data use agreement.

## Consent for publication

Not applicable.

## Funding

This work was supported by NIH R01AG099232 and R01AG080590.

## Author contributions

L.Y., J.R.G., and J.W. designed the research; J.R.G. and J.L.D. provided image annotations; L.Y., J.L.D., and G.M. performed research and analyzed data; L.Y. contributed analytic tools; L.Y., J.R.G., and J.W. wrote the paper; J.L.D., G.M., W.S., G.L., S.U., and H.F. revised the paper; J.R.G. and J.W. jointly supervised the work.

## Competing interests

The authors declare no competing interests.

## Data, code and materials availability

The expert annotations generated in this study are available at https://huggingface.co/datasets/liangyou03/SPInfer-ROSMAP. The ROSMAP cohort data are available through the Rush Alzheimer’s Disease Center Resource Sharing Hub (https://www.radc.rush.edu), subject to a data use agreement. The ROSMAP-Compass snRNA-seq data (Synapse ID syn18485175) are available via the AD Knowledge Portal (https://adknowledgeportal.synapse.org/). The AD Knowledge Portal is a platform for accessing data, analyses, and tools generated by the Accelerating Medicines Partnership (AMP-AD) Target Discovery Program and other National Institute on Aging (NIA)-supported programs to enable open-science practices and accelerate translational learning. The data, analyses and tools are shared early in the research cycle without a publication embargo on secondary use. Data is available for general research use according to the following requirements for data access and data attribution (https://adknowledgeportal.synapse.org/Data%20Access). For access to the data resources described in this manuscript, see https://doi.org/10.7303/9618239. The Visium spatial transcriptomics data were obtained from the publicly available spatialDLPFC study (*9*) and are accessible through the spatialLIBD Bioconductor package (https://bioconductor.org/packages/spatialLIBD). The SPInfer source code, environment specifications, and reproduction workflows are available under the MIT License at https://github.com/liangyou03/SPInfer. The fine-tuned model weights are available at https://huggingface.co/liangyou03/SPInfer-Weights.

## Supplementary materials

### Abbreviations

AD: Alzheimer’s disease
DAPI: 4^′^,6-diamidino-2-phenylindole
FDR: false discovery rate
GATv2: graph attention network version 2
IF: immunofluorescence
IoU: intersection over union
MLP: multilayer perceptron
ROSMAP: Religious Orders Study and Rush Memory and Aging Project
snRNA-seq: single-nucleus RNA sequencing.

**Figure S1:**
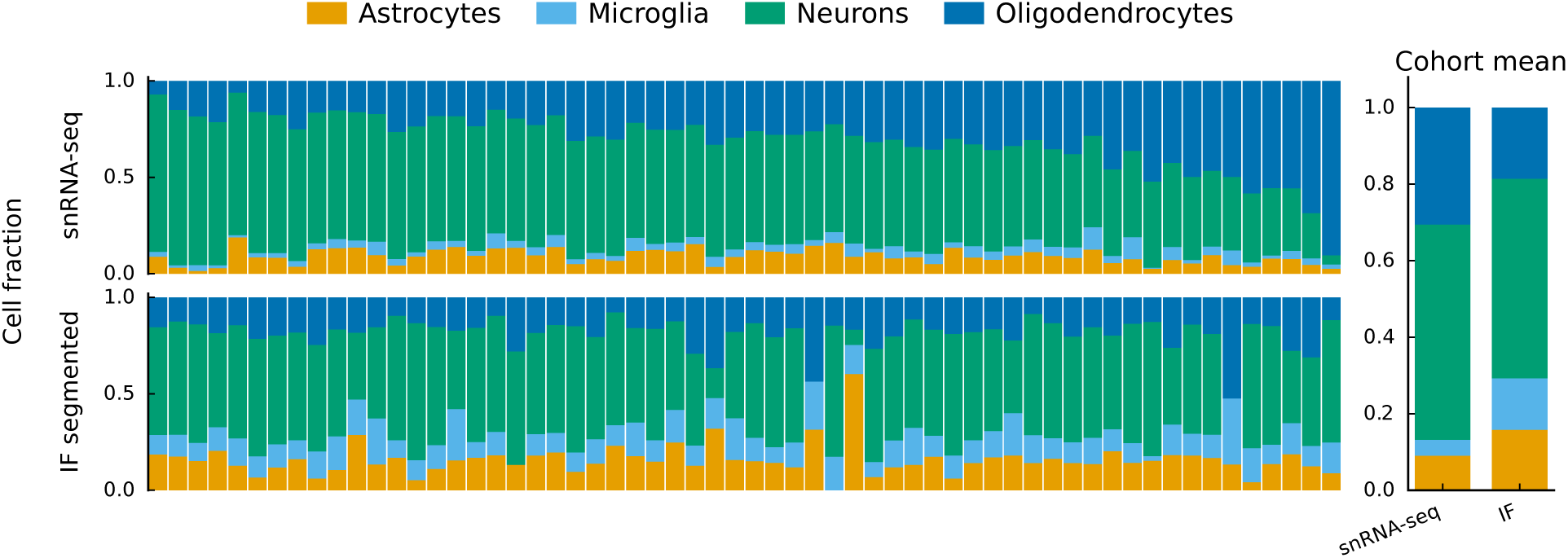
Donor-aligned cell composition profiles in gray matter from snRNA-seq data from the same donors and IF segmentation. Stacked bars show donor-level cell-type fractions for the gray matter single-nucleus RNA-seq data and for IF segmentation-derived estimates, with cohort means shown on the right. Fractions in the single-nucleus data do not sum to 1 because the comparison is restricted to shared cell types and excludes unmatched minor cell types.

**Figure S2:**
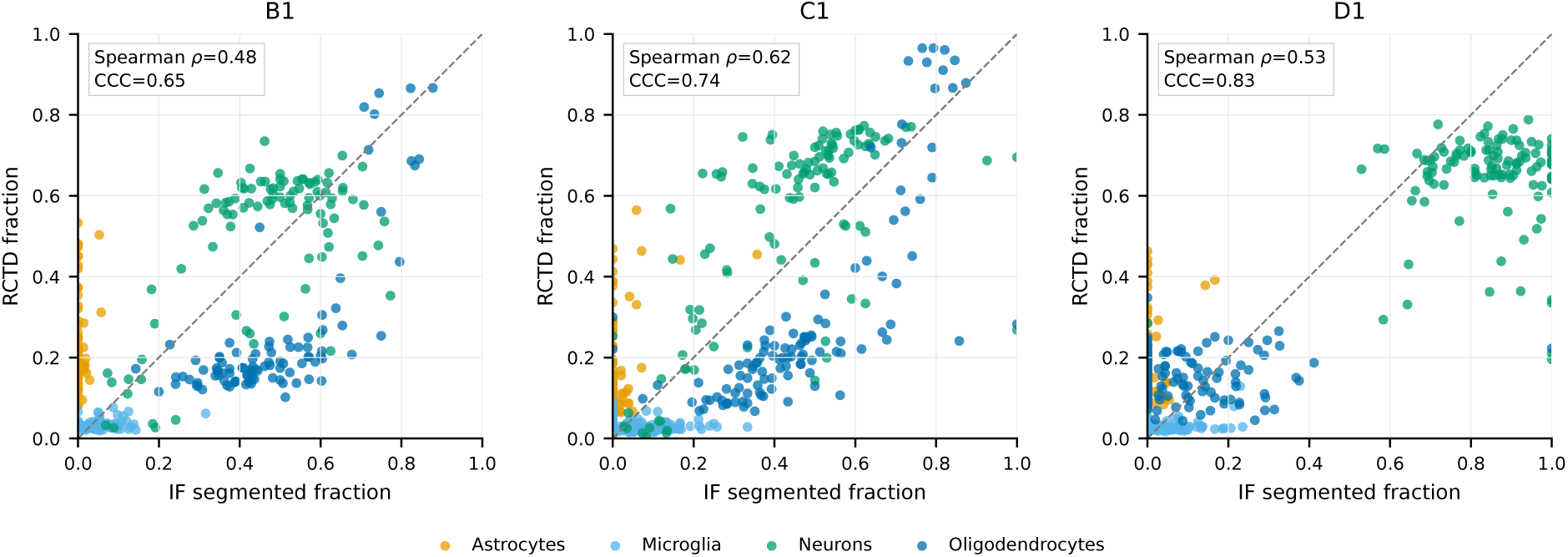
Regional Visium composition agreement in sections B1, C1, and D1. RCTD-derived cell fractions are compared with IF-derived cell fractions after aggregation into spatial bins for the three Visium sections not shown in the main figure. Points are colored by cell type.

**Figure S3:**
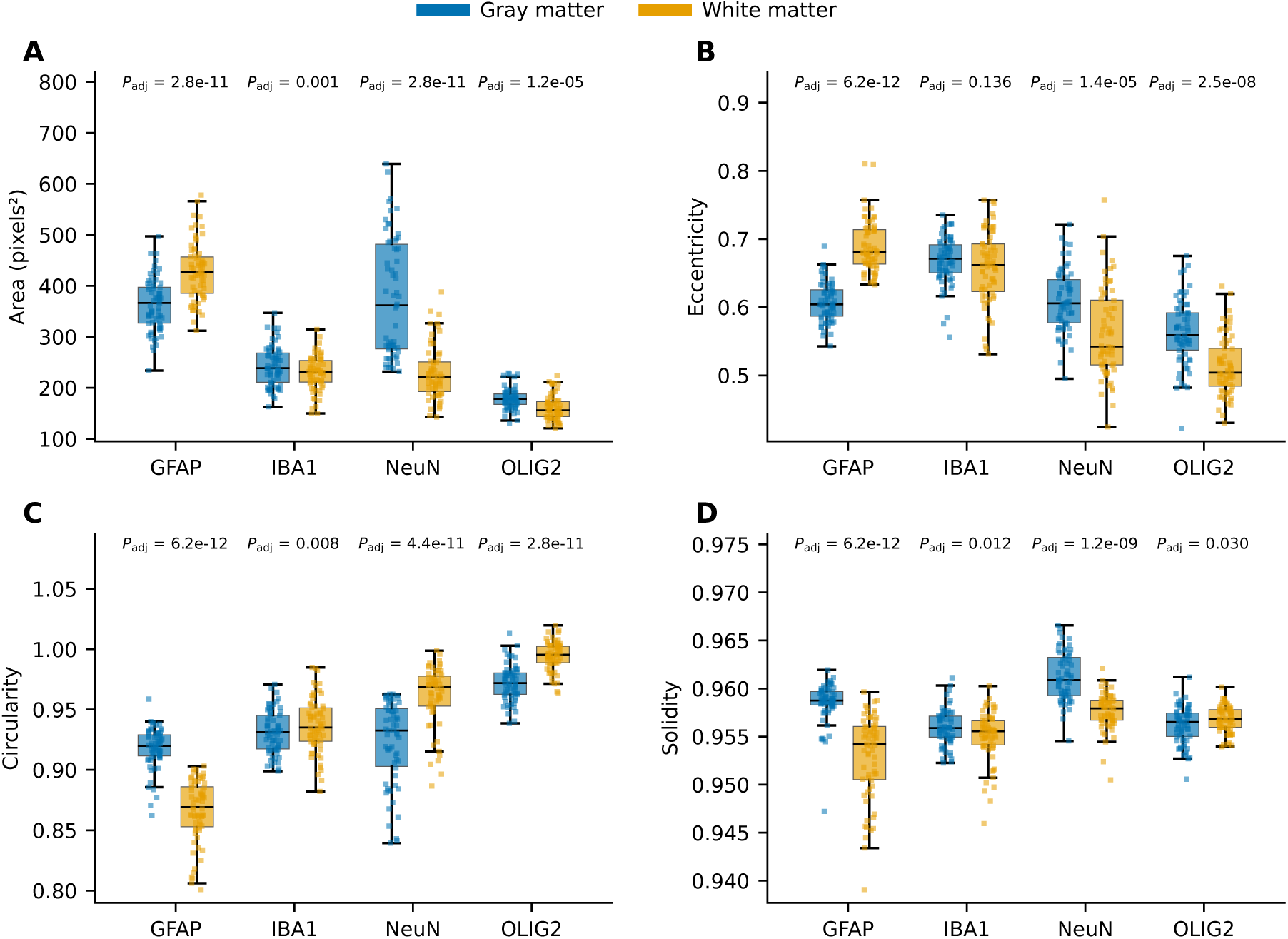
Morphology features by marker and tissue region. Boxplots show donor-level medians of area, eccentricity, circularity, and solidity for marker-positive cells in gray and white matter. Each point represents one donor-region median; only donors with both regions are included. Boxes show the median and interquartile range; whiskers extend to the most extreme values within 1.5 times the interquartile range. Displayed *P*_adj_ values are from two-sided paired Wilcoxon signed-rank tests with Benjamini–Hochberg adjustment across comparisons.

**Figure S4:**
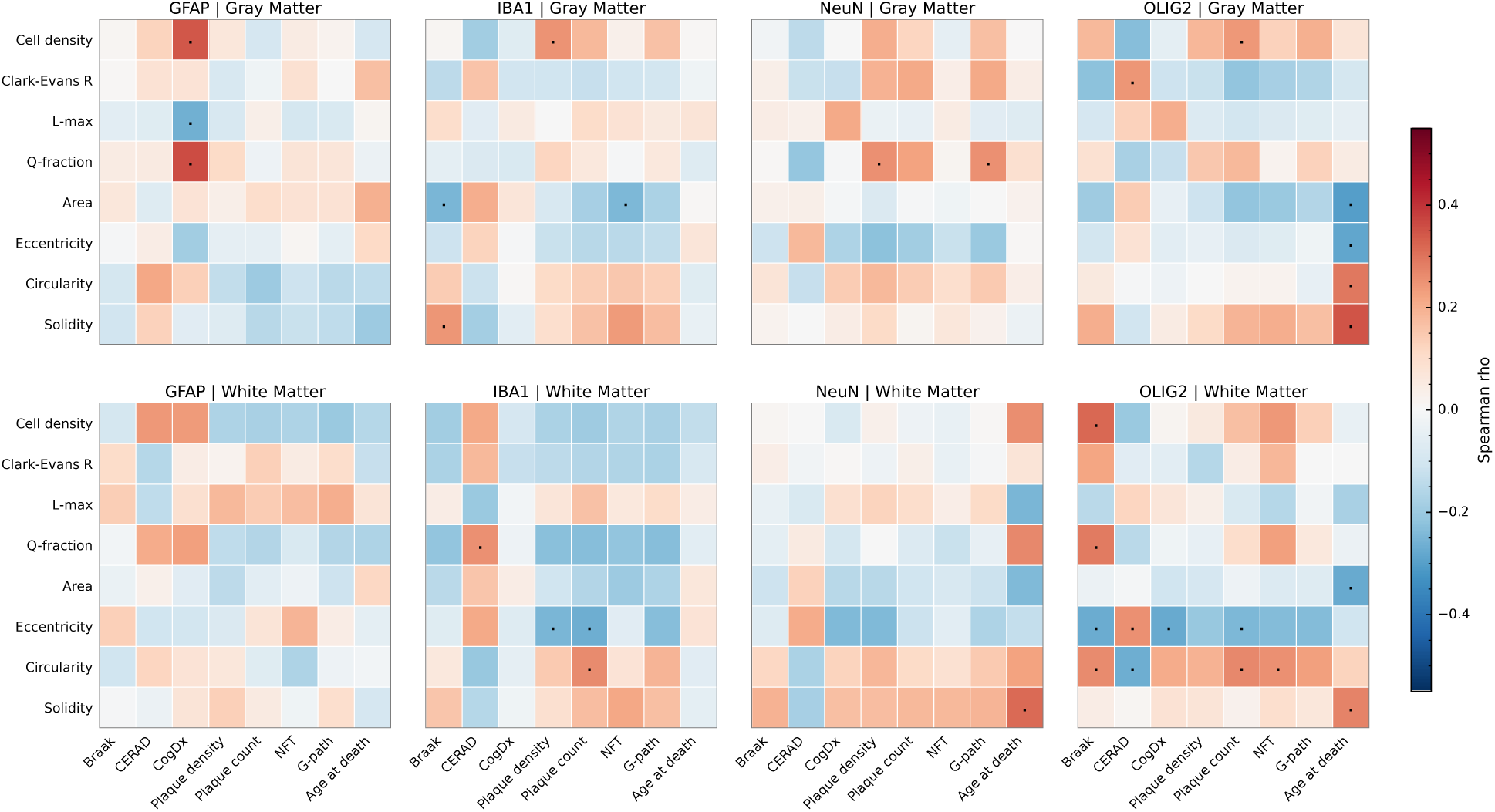
Spearman correlations between spatial or morphological features and clinical variables. Heatmaps show Spearman correlation coefficients for marker- and tissue-specific associations between cell density, spatial organization features, morphological features, and clinical variables. Black points denote nominal *p* < 0.05.

**Figure S5:**
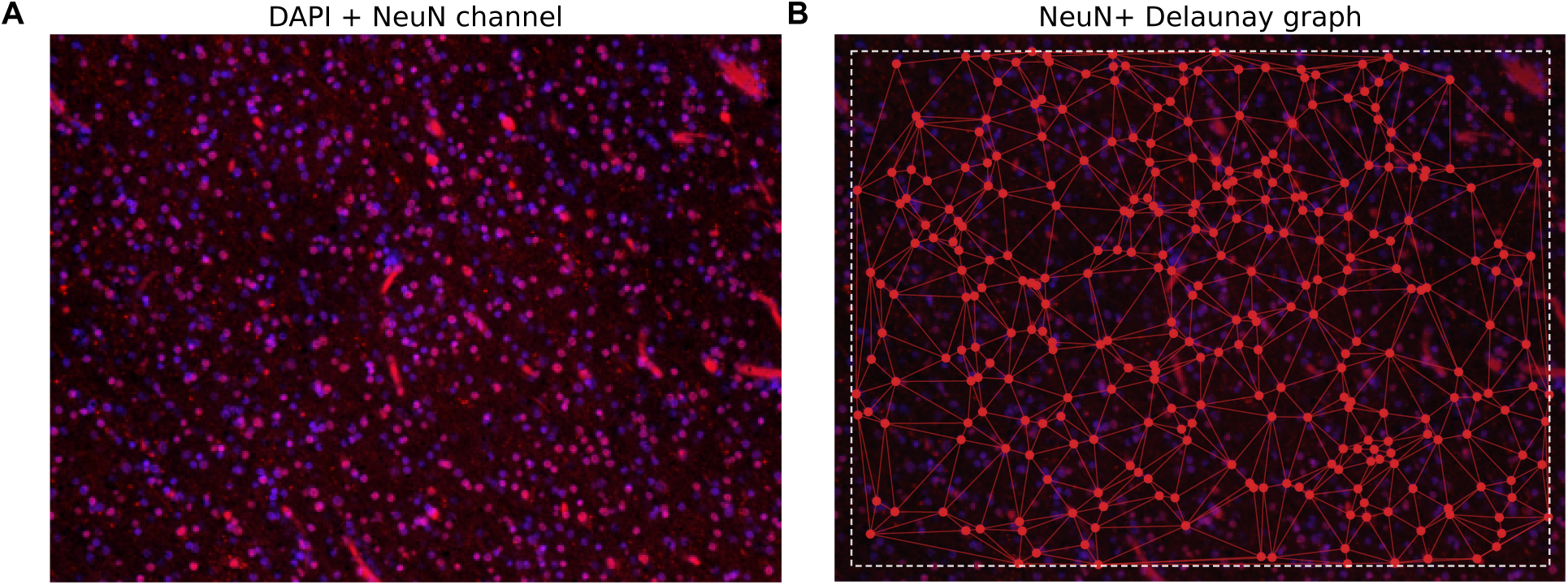
Representative Delaunay graph construction from nuclear centroids within an image, illustrating how bidirectional spatial adjacency edges connect DAPI-positive nuclei prior to graph neural network classification.

**Figure S6:**
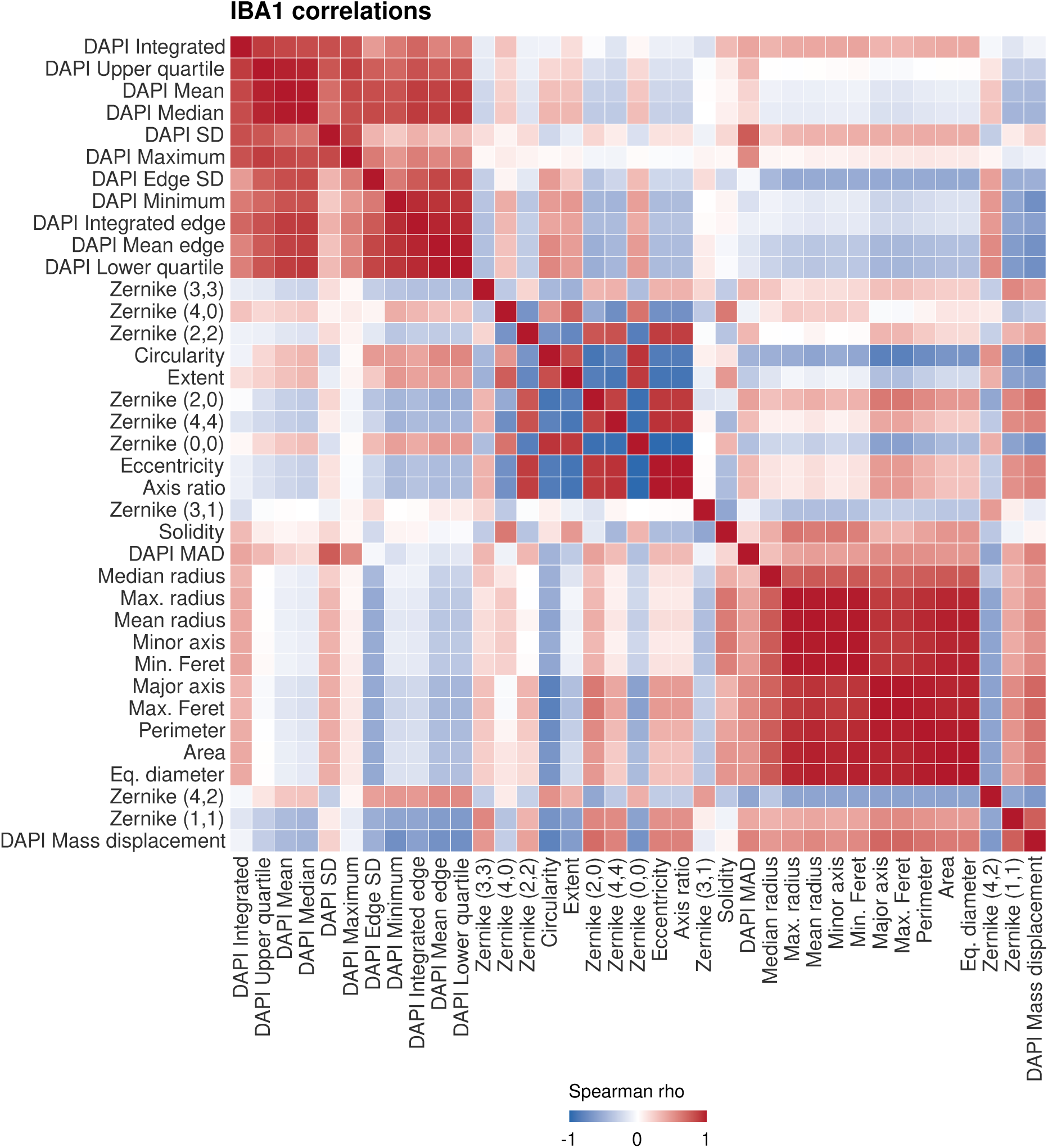
Correlations among all CellProfiler features. Spearman correlations across 56 donors with IBA1-positive cell measurements. Both axes use average-linkage clustering of 1 − |*ρ*|; colors retain the sign of the correlation. All 37 features are displayed.

## Notes

https://huggingface.co/datasets/liangyou03/SPInfer-ROSMAP

https://github.com/liangyou03/SPInfer

### Competing Interest Statement

The authors have declared no competing interest.

### Summary of Updates

Figure 4 has been revised by replacing the background to more accurately reflect the correct data type.

